# Individualised MRI training for paediatric neuroimaging in autism spectrum disorders: A child-focused approach

**DOI:** 10.1101/462234

**Authors:** Emmanuel Peng Kiat Pua, Sarah Barton, Katrina Williams, Jeffrey M Craig, Marc L Seal

## Abstract

Magnetic Resonance Imaging (MRI) in paediatric cohorts is often complicated by reluctance to enter the scanner and head motion-related imaging artefacts. The MRI scanner environment is highly unusual and may distress younger cohorts, especially in children with sensory sensitivity and separation anxiety. The issue is particularly challenging in children with Autism Spectrum Disorders (ASD), where delivery of instructions for novel task demands in an unfamiliar setting may be limited or less effective due to communication disorder or intellectual disability. These issues together often give rise to excessive head motion that can significantly reduce the quality of images acquired, or render data unusable. Here we report an individualised MRI training procedure that enables young children with ASD to better tolerate the MRI scanner environment based on a child-focused approach and an individualised familiarisation strategy. The training procedure was implemented in a locally recruited study of monozygotic twins (n=12; 6 twin pairs; age range 7.1 to 12.85 years) concordant or discordant for ASD. MRI image quality indices were better or comparable to images acquired from an independent multi-centre ASD cohort. The success of these findings is promising and may be useful to improve the quality of paediatric neuroimaging in similar clinical populations.

## Introduction

Autism spectrum disorders (ASD) are a group of lifelong neurodevelopmental conditions characterised by impairments in social communication, and restricted and repetitive behaviours. The reliable identification of atypical brain structure and function in ASD across various modalities using Magnetic Resonance Imaging (MRI) has been a longstanding challenge due to motion-related imaging confounds (<u>Pua, Bowden, & Seal, 2017</u>). MRI is a non-invasive imaging technique for the study of brain structure and function, and has been widely used to understand atypical brain development in neurodevelopmental disorders. However, being in the MRI scanner can be a challenging environment for any child, even more so in the presence of a cognitive or psychiatric comorbidity. During the scanning procedure, individuals are required to remain still for an extended period of time in a foreign, confined and noisy space in isolation. Children are thus likely to be reluctant to enter the scanner, and the resulting discomfort or distress often leads to unwanted head or body movement that can severely impact the quality of imaging data acquired.

Head motion has significant and systematic effects on MRI image acquisition and analyses. The severity and frequency of motion-related artefacts are typically exacerbated in patient compared to control groups, as well as in younger children due to increased movement in the scanner (<u>Van Dijk, Sabuncu, & Buckner, 2012</u>; <u>Yerys et al., 2009</u>). These issues are further compounded in ASD with symptom features such as communication deficits, sensory hypersensitivity, or resistance to change or novel stimuli (<u>Johnson & Myers, 2007</u>). Other comorbidities commonly observed in ASD include attention-deficit/hyperactivity disorder (ADHD) or anxiety, as well as associated communication problems or intellectual disability (<u>Simonoff et al., 2008</u>). Consequently, the increased demand of coping with novel task-demands or significant distress in foreign environments makes successful image acquisition a challenge in this population (<u>Hallowell, Stewart, E Silva, & Ditchfield, 2008</u>).

In recognition of the significant difficulties of neuroimaging young individuals with ASD, there has been increasing efforts to mitigate excessive head motion in ASD and similar at-risk populations using various strategies before and during image acquisition (e.g. <u>de Bie et al., 2010</u>; <u>Greene, Black, & Schlaggar, 2016</u>; <u>Nordahl et al., 2016</u>; <u>Raschle et al., 2012</u>). For example, taking children through a practice or mock MRI session before the actual scan has shown to be an effective intervention for improving the success rate of scanning and improving data quality (<u>Carter, Greer, Gray, & Ware, 2010</u>). There remains a need for specialised strategies to address image acquisition challenges that could be unique across different neurodevelopmental disorders with distinct clinical presentations, such as in ASD. Building on current knowledge and previous recommendations, we designed a training protocol to prepare young children with ASD for MRI neuroimaging in consideration of symptom features of the condition. The primary goal was to enable participants with ASD to better tolerate MRI imaging with acceptable levels of head motion. Here we report the implemented training protocol and quality of MRI image acquisition for a locally recruited monozygotic twin study on ASD. Framewise displacement (FD) as an estimate of change in head motion across image volumes that strongly relates to motion artefacts, and rate of change of acquired signal across the whole brain (DVARS), were used as data quality indexes to compare image quality of the locally recruited sample to a large multi-centre ASD cohort.

## Method

### Participants

Participants (n=12) were locally recruited from Twins Research Australia (TRA) and an ASD epigenetics study at The Royal Children’s Hospital (RCH; HREC 33208C). Inclusion criteria were monozygotic twins concordant or discordant for ASD between the ages of 5 to 18 years, of either sex, and raised in the same household in the greater metropolitan area of Melbourne, Victoria. The lower limit of the age range was selected due to the known challenges of scanning tolerance and motion artefacts in imaging young children, especially in atypical neurodevelopmental populations. ASD diagnosis was previously determined by clinical assessment on the gold standard Autism Diagnostic Observation Schedule-2 (ADOS-2) or by the TRA with supporting medical documentation of prior diagnosis and assessment. Zygosity status of twin pairs was confirmed with genetic testing using a twelve-marker panel following DNA extraction from buccal swabs. Results of zygosity testing were only released to parents upon request to respect the privacy of families. Informed consent was obtained from all participants and the research study protocol was approved by the RCH Human Research and Ethics Committee (HREC 36124C). All research was performed in accordance with the Code of Ethics of the World Medical Association (Declaration of Helsinki). Table 1 provides a descriptive summary of participant demographics.

**Table 1.** Participant demographics and data quality indexes

| Twin Pair | Gender | Age<br>(years) | ASD<br>Diagnosis | SRS | FD<br>(Run 1) | DVARs<br>(Run 1) | FD<br>(Run 2) | DVARs<br>(Run 2) |
| --- | --- | --- | --- | --- | --- | --- | --- | --- |
| 1 | Male | 10.43 | Yes | 51 | 0.20 | 1.16 | 0.40 | 1.14 |
| 1 | Male | 10.43 | Yes | 101 | 0.13 | 1.20 | 0.25 | 1.19 |
| 2 | Male | 9.82 | Yes | 116 | 0.87 | 1.14 | 0.48 | 1.16 |
| 2 | Male | 9.82 | Yes | 136 | 0.31 | 1.17 | 0.35 | 1.25 |
| 3 | Female | 7.1 | Yes | 145 | 0.16 | 1.27 | 0.15 | 1.35 |
| 3 | Female | 7.1 | Yes | 124 | 0.13 | 1.27 | 0.13 | 1.26 |
| 4 | Male | 12.85 | Yes | 139 | 0.14 | 1.16 | 0.18 | 1.19 |
| 4 | Male | 12.85 | No | 109 | 0.30 | 1.18 | 0.30 | 1.12 |
| 5 | Male | 12.17 | Yes | 122 | 0.19 | 1.20 | 0.22 | 1.21 |
| 5 | Male | 12.17 | No | 49 | 0.16 | 1.15 | 0.15 | 1.18 |
| 6 | Female | 7.49 | Yes | 100 | 0.19 | 1.16 | 0.18 | 1.12 |
| 6 | Female | 7.49 | No | 7 | 0.21 | 1.22 | 0.22 | 1.20 |

*Note.* SRS: Social Responsiveness Scale-2 total score; FD: Framewise displacement; DVARs: standardized DVARs measure (rate of change of image intensity relative to the previous timepoint; Powers 2012).

### Pre-visit preparation

Standard MRI visit protocols were modified for each participant to accommodate expected difficulties specific to ASD. Key goals were to familiarise participants to the MRI scanner environment, and to improve tolerance to the loud and repetitive acoustic noise in the scanner. First, parents of children that met the eligibility criteria were contacted for a pre-visit interview around one to two weeks before the MRI visit. A brief semi-structured clinical interview was conducted by a board-certified provisional psychologist, with the aim of gathering relevant information to develop reinforcement and motivation strategies individually tailored to each child. The interview included queries about general interests and hobbies, and reward strategies that were effective for that particular child. The goal of such strategies was to increase the likelihood of desirable behaviours with positive reinforcement in the form of verbal or material rewards. The approach has previously shown to be effective for behavioural modification outcomes in ASD (<u>Gena, Couloura, & Kymissis, 2005</u>). Triggers that typically preceded idiosyncratic episodes of distress or repetitive symptoms were also identified for children with an ASD diagnosis. Potential strategies for intervention were explored to better understand the parent-child interaction during such episodes, and possible approaches to manage episodes that may occur on-site during the MRI visit.

Parents were then provided with an in-house MRI familiarisation package that comprised of three main components:

1. A brief, two to three minutes duration, MRI orientation video^1^ filmed at the RCH introduces the child to locations in the hospital that would be encountered during the actual on-site visit, such as the patient waiting area, preparation room, and MRI scanner.
2. An ‘Okee in Medical Imaging’ application^2^ developed by the RCH. This free-of-charge application can be easily accessed on most smart device platforms, and contains a suite of games designed to familiarise young children with aspects of neuroimaging, as well as information and tips for parents to help their child prepare for the visit. The training games suitable for MRI preparation include introducing the child to different parts of the MRI scanner in an undersea submarine pretend-play simulation, and adventure games to teach a child how to keep still when a signal is given. The aim of these games was to help the child understand concept of minimizing in-scanner movement across different intervals, and to engage in the appropriate behaviour when instructed. The Okee application also contains similar games designed for other common medical imaging modalities such as CT, ultrasound, and X-ray.
3. Parents were also provided with recorded audio clips of sounds emitted by the MRI scanner that each child would later experience during the actual scan (see Supplementary material). The acoustic noise is generated by vibrations of the MRI gradient coil, and its cyclic repetitive pattern or loud volume may lead to discomfort resulting in an undesirable increase in head or body motion (<u>Cho et al., 1997</u>; <u>Counter, Olofsson, Grahn, & Borg, 1997</u>). The audio clips in the familiarisation package comprised of the actual acoustic noise generated by the specific MRI sequence used in the research study. The samples were recorded in real-time from our scanner and thus extracted by our imaging technicians. This is necessary because generated acoustic noise could vary between scanner sites, equipment, and acquisition sequences. Parents were given instructions to familiarise their children to the MRI sound clips on a daily basis, beginning with a brief playback of the audio clip on the initial day followed by a gradual increase in the length of playback over a one-week period. Any type and level of distress elicited by the audio clips were to be noted and reported. Parents were also asked to show their children the MRI orientation video at least once, and to allow them to explore the Okee medical imaging application. Parents were encouraged to bring comfort objects to reduce anxiety in the child, and a favourite movie for the visit. Links to the familiarisation package are provided in footnotes or Supplementary materials.

### Hospital visit

On the day of hospital attendance, the visit comprised of two key phases: a mock MRI training session and the actual MRI scan. Based on information gathered during the pre-visit interview, both the mock and actual MRI were personalised to suit individual needs and language ability of each participant For example, if parents noted that their child tended to become distressed with unfamiliar people or situations, rapport building would be a key focus during the initial contact and period of interaction with families. Nonverbal approaches and materials were emphasised for participants with poorer language ability. Staff identification cards were also used to facilitate the learning of name-face associations based on explicit audio-visual cues, rather than a generic or brief introduction by name.

### MRI orientation session

The MRI introduction session comprised of three phases; an explanation of the day’s schedule (4-5 hours), a play-based session explaining the MRI (30 minutes) and interacting with the mock MRI session (30 minutes). The visit began with an outline of the planned activities for the day. Given that an established and predictable schedule is often useful to facilitate the completion of tasks in individuals with ASD (<u>Horner, Carr, Strain, Todd, & Reed, 2002</u>), the aspect of time was clearly emphasised, to the point of providing specific start and end times for each activity if the information was deemed helpful for the child. A visit map of the appointment schedule was prepared as a visual aid in the form of a flowchart with photographs of the actual location or item associated with a particular activity for each task:

1. Introduction (photograph: hospital common waiting area)
2. Visit orientation (photograph: mock interview room)
3. Mock training session (photographs: MRI mock scanner, DVD library of movies)
4. MRI waiting area (photograph: MRI reception desk)
5. MRI scanning session (photograph: MRI scanner)

Photographs of key locations or equipment in the flowchart were used as visual cues to familiarise the child with each segment of the visit. The MRI orientation video from the familiarity package was presented again to each participant, and they were encouraged to explore the Okee application. Participants were then informed about rewards they would receive at the end of the visit. These rewards were personalised for each individual based on motivation strategies discussed with parents during the pre-visit interview and served as a form of positive reinforcement. The items were specific to each child and ranged from art materials to trading cards or a soft toy. Each participant further received a certificate for completing the mock training, a certificate for completing the MRI scan, and a screenshot of their scan as a keepsake.

Next, the child was engaged in a play-based session to explain the MRI scanning process. A pictorial storyboard comprising of photos of on-site facilities and equipment was used to describe the functioning of a MRI scan in simple terms. The importance of keeping still was heavily emphasised, and pictures of scans with excessive motion artefacts were shown to demonstrate the effect of head motion on image quality. The acoustic noise experienced during the scan was explained to be similar to the audio clips the child would have been listening to on a regular basis in the week prior to the visit. A cartoon illustrated storybook incorporating various elements of the MRI scan procedure in the form of a social story was available for the same purpose as well. After the storyboard presentation, the scanning process was recreated using a pretend play-set with dolls and customized wooden blocks for the child to observe. The playset was used to sequentially explain what a participant might experience during a typical scan, and to introduce different components of the scanner such as the horizontal and vertical movement of the MRI bed and the helmet head coil.

### Mock MRI simulation

In the subsequent mock MRI simulation phase, participants were given the opportunity to interact with a non-functional MRI scanner. The mock scanner is equipped with MRI scanner components that the child would typically interact with during the actual scan, such as the moveable patient bed, head coil, and headphones. Participants were systematically introduced to the scanning process using a task-analysis approach, in which a task is segmented into a sequence of smaller steps or activities (<u>Hernandez & Ikkanda, 2011</u>). This allows the participant to familiarise themselves with the scanner environment at their own pace, and to reduce the risk of the child becoming overwhelmed. The stepwise approach offers the opportunity for additional instruction or support at each step should a participant be showing signs of distress or anxiety, and progression to the next stage only occurs once a child has comfortably completed the previous step. Stages for the mock scanning process were as follows:

1. Entering the mock scanner room
2. Exploring the mock scanner
3. Playing a DVD movie of their choice (from home or available library)
4. Approaching the mock scanner bed
5. Operating the mock scanner bed (vertical and horizontal movement controls)
6. Sitting on the mock scanner bed
7. Putting on headphones
8. Lying on the bed
9. Tolerating vertical movement of mock scanner bed
10. Placement of the head coil
11. Listening to MRI gradient noise outside the scanner (external playback of recorded MRI audio clips from familiarisation package)
12. Lying still on mock scanner bed for one minute
13. Tolerating horizontal movement of mock scanner bed into scanner
14. Listening to the MRI gradient noise in scanner
15. Increasing amount of time to lie still (5 minutes) while listening to MRI audio clip in the scanner and watching a movie

The stepwise approach was used to gradually familiarise each participant with the scanning process at their own pace and comfort levels. Verbal encouragement or material rewards was used to reinforce successful completion of each step where necessary. If participants brought along a comfort object, they were allowed to hold on to it when entering the mock scanner. If the object was a figure or soft toy, it was also used as a mock participant to demonstrate different steps of the scanning process. Importantly, every participant were also allowed to observe their co-twin undergo the mock simulation process to facilitate learning through peer-modelling and sibling involvement (<u>Shivers & Plavnick, 2014</u>).

To estimate participant head motion in the mock scanner, measurements from an accelerometer device were recorded in the final step where the child was instructed to remain still for five minutes on the scanner bed while listening to a playback of MRI gradient noise, and watching a movie. The accelerometer device (3-axis, 50 Hz sample rate, 15-bit resolution) was attached to the mock scanner headphones and wired to a computer terminal in the same room, providing real-time feedback of participant motion in the mock scanner simulation. The co-twin of participants observed their sibling for the full duration of the mock training session, before undergoing the training themselves as their sibling observed. Using a real-time display of signal from the accelerometer device, participants were also given visual feedback to appreciate how excessive head motion introduces noise and visible fluctuations in the signal being recorded. As the goal of mock scanner simulation was to facilitate successful and comfortable completion of a simulated MRI scan (i.e. the final stage) with minimal movement, the suitability of each participant for an actual scan was assessed at the end of the training session. Participants were debriefed to explore and address any further concerns from participants or their parents. Families were allowed to take pictures with the mock scanner and each child was rewarded with a certificate of completion for the mock training phase. Participants were given a one to two hour break before their MRI scan.

### MRI Acquisition

Following a recently published MRI protocol specifically designed for neurodevelopmental investigations (<u>Silk et al., 2016</u>), multimodal MRI data was acquired on a 3-Tesla Siemens TIM Trio MRI scanner (Siemens, Erlangen, Germany) with a 32-channel head coil. To estimate measures of brain structural morphometry (e.g. cortical thickness, surface area), a modified multi-echo magnetization prepared rapid gradient-echo (MEMPRAGE) sequence was used to acquire T1-weighted anatomical images (TR=2530ms, TE=1.77,3.51,5.32,7.20ms, TI=1260ms, flip angle=7.0 deg, voxel size=0.9 x 0.9 x 0.9 mm, FOV read=230mm). Navigator based prospective motion correction was implemented with Siemens in-scanner motion correction (MoCo), where field-of-view and slice positioning is updated to adjust for motion in real-time to reduce motion artefacts and improve image quality. Participants were allowed to watch a movie of their choice during image acquisition.

Task-free blood oxygen level-dependent (BOLD) fMRI data to estimate functional connectivity between brain regions based on intrinsic correlated neural activity at rest was acquired with multi-band accelerated EPI sequences across two separate runs within the same scanning session (TR=1500ms, TE=33ms, volumes =250 voxel size = 2.5 x 2.5 x 2.5mm, multi-band factor=3). Participants were instructed to keep their eyes open and attention focused on a fixation cross during the scan.

Diffusion-weighted imaging (DWI) data to investigate white-matter microstructure was also acquired with multi-band accelerated EPI sequences for multi-shell acquisition (b=2800, 2000, 1000 s/mm^2^ + interleaved b=0 s/mm^2^, phase encoding direction: anterior to posterior). Standard and reverse phase encoded blipped images with no diffusion weighting were obtained to correct for magnetic susceptibility-induced distortion.

The overall scan duration for each subject was around 45 minutes, with brief periods of rest between sequences where participants were allowed to move and stretch. All participants successfully completed the scans without withdrawing or displaying significant signs of distress. Each child was rewarded with another certificate of completion, along with their individual rewards as described above.

### Data quality analyses

Framewise displacement and DVARS were used as quality control metrics from the MRI Quality Control Tool (<u>MRIQC; Esteban et al., 2017</u>). The tool integrates modular sub-workflows dependent on common neuroimaging software toolboxes such as FSL, ANTs and AFNI. The data was first minimally preprocessed in the MRIQC anatomical workflow with skull-stripping, head mask and air mask calculation, spatial normalization to MNI space, and brain tissue segmentation. T1-weighted images were visually inspected for ringing artefacts, blurred grey- and white-matter boundaries, and background noise (<u>Pardoe, Hiess, & Kuzniecky, 2016</u>). Head motion correction was performed in the functional workflow with AFNI *3dvolreg* (<u>Cox, 1996</u>). The algorithm computes head realignment across frames, registering each image to a base reference volume using a six-parameter, rigid-body transformation (angular rotation and translation). Framewise displacement (FD) is a data quality index expressing instantaneous head-motion based on change in head position across frames. FD is the estimated spatial deviation between the reference volume and all other volumes derived from sum of the absolute values of the differentiated rigid-body realignment estimates. Rotational displacements were computed as the displacement on the surface of radius 50mm. Framewise displacement was additionally with a different tool (FSL *mcflirt)* for validation (<u>Jenkinson, Bannister, Brady, & Smith, 2002</u>). DVARS is another quality index that estimates the rate of change of BOLD signal across the whole brain at each frame (<u>Smyser et al., 2010</u>). The change in image intensity compared to the previous timepoint was computed by differentiating the volumetric timeseries and obtaining the root-mean-square signal change. The metric was normalized with the standard deviation of the temporal difference timeseries to allow comparisons between different imaging sites and scanners (<u>Nichols, 2017</u>).

For multi-centre and multi-cohort comparisons, data quality metrics were obtained from the Autism Brain Imaging Data Exchange (<u>ABIDE-II; Di Martino et al., 2017</u>). ABIDE-II is a publicly available aggregation datasets from n=487 individuals with ASD and n=557 controls (age range: 5-64 years). Site-specific protocols for participant preparation and image acquisition are available^3^. Quality metrics of ABIDE-II data was previously derived using the Quality Assessment Protocol (QAP) from the Preprocessed Connectomes Project (<u>Shehzad et al., 2015</u>).

## Results

All participants underwent each stage of the MRI training protocol (pre-visit phase, orientation session, mock MRI simulation and evaluation). Parents provided verbal confirmation of participant engagement with the familiarisation package. Materials from each component were also presented to participants again during the orientation session. The entire duration of the hospital visit ranged from four to five hours per family, including a one-hour lunch break. Mean FD across the locally recruited sample representing instantaneous head motion for both task-free sequences (Run 1: *M=* 0.25 mm, *SD=*0.20; Run 2: *M=* 0.25 mm, *SD=*0.11) were more favourable compared to individuals with ASD from the ABIDE-II cohort (*M=* 0.31 mm, *SD=*2.37), but not typical controls (*M=* 0.11 mm, *SD=*0.14). FD results obtained from FSL *mcflirt* (Run 1: *M=* 0.23 mm, *SD=*0.19; Run 2: *M=* 0.23 mm, *SD=*0.10) were comparable to the output from *AFNI 3Dvolreg,* suggesting stability of FD estimates independent of analysis method. DVARS as the standardized root mean squared change in BOLD signal intensity from one volume to the next in the local sample for both runs (Run 1: 1.19; Run 2: 1.20) were similar to individuals from the ABIDE-II cohort (ASD: 1.16; controls: 1.16). Only one participant from the local cohort failed to meet the threshold of <0.5mm for acceptable FD due to excessive head motion (95.8% success rate). The same participant however demonstrated improved FD (0.48mm) below the threshold in the repeat run later acquired within the same scanning session, a significant reduction from the observed FD in the initial run (0.87mm).

Based on monozygotic twin regression modelling from <u>Carlin, Gurrin, Sterne, Morley, and Dwyer (2005)</u>, within-twin-pair differences in outcome variables were regressed onto within-pair differences in independent variables. Intrapair associations between FD from the initial and repeat resting-state acquisition was near but not significant (t=2.386, *R*^*2*^*=*0.44, *p=*0.06). As expected, FD was strongly related to DVARS (Run 1: t=3.79, *R*^*2*^*=*0.69, *p=*0.01) suggesting a relationship between head motion and BOLD signal intensity change between volumes. The association was present but less robust in the repeat acquisition (Run 2: t=2.24, *R*^*2*^*=*0.40, *p*=0.07). Vertical head motion recorded during the mock training simulation predicted in-scanner head motion during the first resting-state acquisition (Run 1: t=-2.52, *R*^*2*^*=*0.47, *p=*0.05; Run 2: t=-0.81, *p=*0.45; Z-axis root-mean-squared-successive-difference). There were no within-twin-pair associations for FD and DVARS in the initial (FD: *r=*0.57, *p*=0.24; DVARS: (*r=*0.34, *p*=0.51) and repeat acquisition (FD: *r=*0.57, *p*=0.24; DVARS: *r=-*0.56, *p*=0.25).

## Discussion

This paper presents an MRI training procedure for participants with or without ASD in a paediatric twin cohort. Key components were a personalised child-centered approach and an MRI familiarisation strategy. Briefly, a child-focused strategy was developed for each individual participant based on a pre-visit clinical interview with parents to identify effective methods for motivation and potential distress or symptom triggers. Families participated in an MRI familiarisation procedure one week prior to the visit, during which participants were gradually introduced to MRI acoustic gradient noise on a daily basis. Parents were also provided with materials such as an MRI orientation video and games application to familiarise their children with the MRI scanner, and were encouraged to engage in the activities together. On the actual day of the visit, a significant proportion of time was allocated to each stage of the MRI training protocol. Key components were an orientation phase with the use of visual aids and previously distributed familiarisation materials, a play-based session incorporating an MRI playset, and a task-based approach was used to simulate the scan procedure in a mock MRI scanner before the actual scan. Importantly, implementation of the process was flexible and readily adapted based on information gathered from the pre-visit interview to best suit the needs and ability of each participant (see Methods for full details; Figure 1).

**Figure 1.**
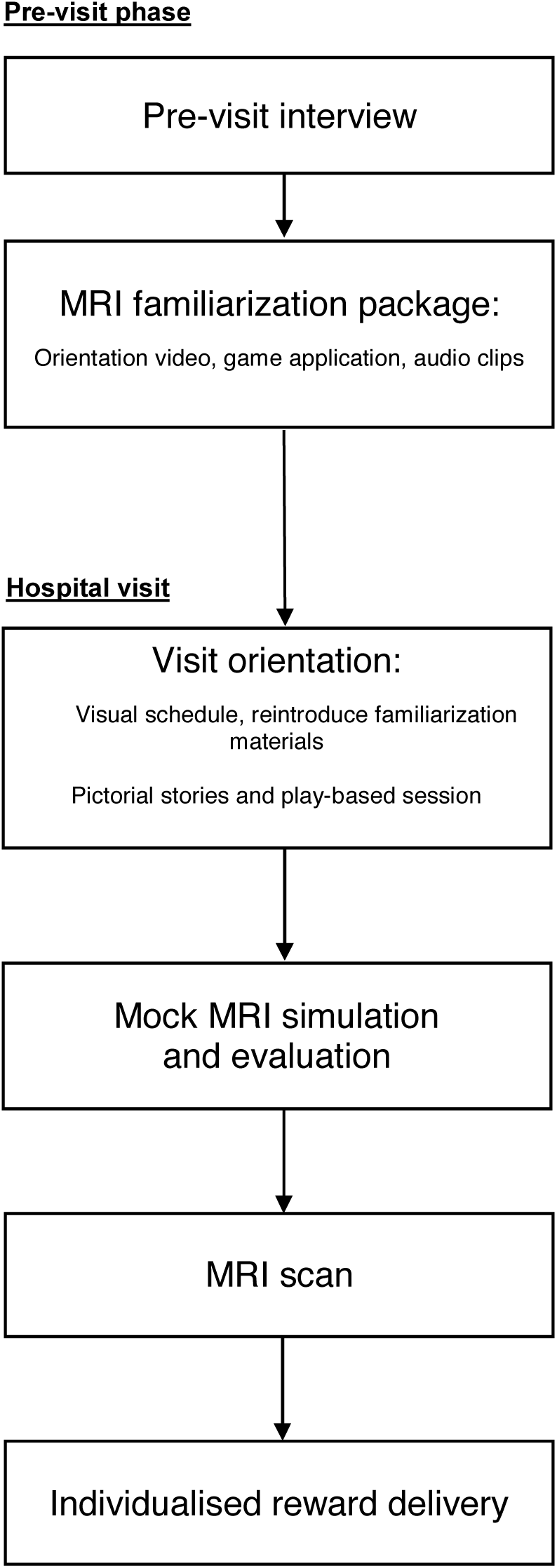
Flowchart of MRI participant training protocol.

Based on the thresholds for acceptable levels of head motion and image artefact quality control, only one participant failed to meet criteria (95.8% success rate). Mean head motion was significantly reduced in repeat image acquisition within the same session for the same participant. Across all participants, head motion in the initial acquisition was not significantly associated with that observed in the later run. Overall, this suggests that repeat acquisition sequences may be an effective tempering strategy for individuals with excessive head motion. The utility of repeating sequences of interest is in agreement with previous recommendations to increase power and volumes retained after motion correction and denoising procedures in similar paediatric cohorts (<u>Greene et al., 2016</u>). Importantly, image quality indexes for head motion and signal change in the present study exceeded or was similar to the quality of data from the multi-centre ABIDE-II cohort for individuals with ASD (Figure 2). As expected, individuals with ASD from across cohorts displayed more head motion compared to neuroptypical controls. Given the difficulties of neuroimaging paediatric cohorts, data quality outcomes based on these findings suggest that the MRI training procedure may be useful in mitigating motion-related artefacts.

**Figure 2.**
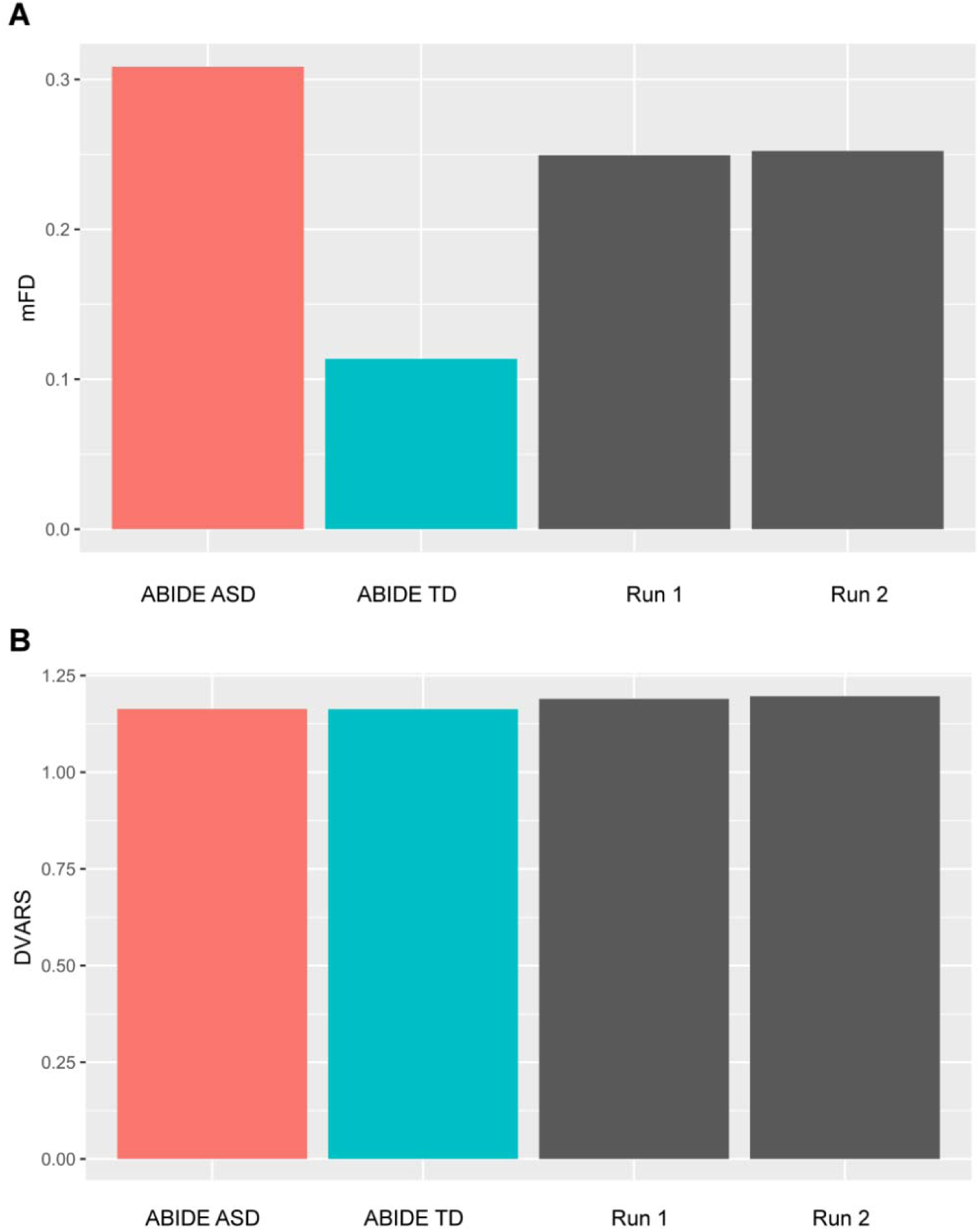
Image data quality indexes with comparisons to the ABIDE-II cohort. mFD: Framewise displacement. Higher values indicate increased volume-to-volume head motion; DVARS: Standardized DVARS measure. Higher values indicate increased change in image intensity across volumes; ABIDE ASD: ABIDE-II individuals with ASD; ABIDE TD: ABIDE-II neurotypical controls; Run 1: Initial task-free sequence for local sample; Run 2: Repeated task-free sequence for local sample.

### A child-focused approach

The common adage that no two individuals on the ASD spectrum are alike underpins the understanding that each individual child with ASD is unique in how core symptoms present due to the complex phenotypic heterogeneity of the condition (<u>Masi, DeMayo, Glozier, & Guastella, 2017</u>). The benefits of child-focused interventions in ASD based on individualised techniques to target specific behavioural outcomes or symptom reductions are similarly well established (<u>Tonge, Bull, Brereton, & Wilson, 2014</u>). The MRI training protocol implemented here was designed to be personalised and tailored to the individual participant. Information gathered in the pre-visit interview critically informs strategies for the mock MRI training session during the visit, where different components of the protocol are emphasised or adapted based on the child’s needs or level of functioning.

The pre-visit interview facilitates information gathering that is critical to the planning and delivery of the training protocol for each participant. Functional assessment was previously reported to be the most consistent factor in predicting intervention success, where effectiveness appears to increase with the precision of assessment in some form of interview, direct observation or functional analysis (<u>Horner et al., 2002</u>). In particular, informant information can provide a comprehensive picture of behaviours and symptoms of the individual child across multiple environments and settings (<u>Stratis & Lecavalier, 2014</u>). In the context of the present study, parent information derived from the pre-visit interview facilitated the development of a personalised strategy for each child during the mock training phase. Individualised information about each child critically supports the later training phase that incorporates several elements of Positive Behaviour Support (PBS) interventions, such as removing triggers or antecedent events preceding undesirable behaviours, teaching new skills, and rewarding positive behaviours (<u>Carr et al., 2002</u>). PBS outcomes have been shown to improve further when informant information across different contexts was available, in addition to partnership efforts with multiple parties including the family and the school (<u>Harvey, Lewis-Palmer, Horner, & Sugai, 2003</u>).

By understanding the interests, motivations, and fears of each child during the pre-visit interview, personalised and effective positive reinforcement strategies were developed together with parents in a collaborative partnership. Using this approach, we found that participants were more likely to comply with instructions and better tolerate the MRI preparation and scanning procedures. Given that social or verbal praise and attention may not be as rewarding for some children with ASD, the identification and implementation of individually functional reinforcers is an important component of the strategy. For example, personalised rewards tailored to the interests of each child (e.g. favourite snack or toy) could serve as more effective positive reinforcers (<u>Horner et al., 2002</u>). Essentially, child-focused approaches to meet task-demands are based on adaptable strategies to uniquely support the needs of each individual child and family (<u>Trivette, Dunst, & Sandall, 2000</u>). Throughout the entire process, parents and participants were reassured that their child’s well-being took priority over research outcomes, and were allowed to withdraw at any stage during the training or scanning phase of the visit.

### Familiarisation strategy

The other key aspect of the training protocol was the incorporation of familiarisation strategies for the MRI scanner environment, with participant tolerance to the loud and repetitive acoustic noise in the scanner as a specific target. Drawing from basic elements of graduated exposure therapy (<u>Craske, Treanor, Conway, Zbozinek, & Vervliet, 2014</u>; <u>Wolpe, 1968</u>), participants were systematically exposed to variations of the scanner environment that progressively became closer approximations to the actual scan. Graduated exposure techniques have been shown to be effective interventions to overcome setting avoidance or reduce anxiety in high- and low-functioning individuals with ASD (<u>Hagopian & Jennett, 2008</u>). Similar to our observations, the combination of graduated exposure and positive reinforcement to reduce setting and activity avoidance was particularly effective (<u>Schmidt, Luiselli, Rue, & Whalley, 2013</u>). Initial exposure to the scanner environment began with the audio playback of the acoustic noise in their homes one week prior to the hospital visit, followed by the RCH-specific orientation video and game that visually introduced the child to the actual scanner location in the hospital, and finally the physical introduction to the scanner environment and the graduated mock scanner simulation leading up to the actual MRI scan. Progression was based on successful completion of the previous stage with minimal anxiety or discomfort, and the speed of progression was adjusted depending on the performance of each participant where applicable. The goal was to facilitate desensitization through gradual exposure to the target stimulus, allowing participants to develop tolerance or habituate to in-scanner noise over time progressively and to minimize anxiety-related or avoidant responses. A flowchart of on-site photographs mapping the schedule and length of activities was additionally used to visually structure the visit for participants (see Supplementary materials). The implementation and utility of a visual schedule is similar to the Picture Exchange Communication System (PECS) in which instruction or learning is supported by visual elements that complements or minimizes verbal input. The strategy offers the child a consistent and predictable system to understand a sequence of activities or tasks, and has been highly effective in facilitating communication and instruction in this population (<u>Schneider & Goldstein, 2010</u>; <u>Shane, 2006</u>).

Another important feature of the familiarisation procedure was high family-involvement. The initial delivery of the MRI familiarisation materials by a parent in the home environment is supported by previous reports showing that efficacy of ASD interventions improved when method of delivery included familiar agents (e.g. parents or teachers) in typical contexts (<u>e.g. home or school; Horner et al., 2002</u>). As children with ASD may have difficulty applying learned skills or outcomes in a novel settings, interventions or learning strategies should be initially initiated by a familiar person and integrated with a child’s daily routine and activities in a natural learning environment (<u>Childress, 2004</u>). This is an important lead-up to the mock MRI on the actual day of visit, where parents are encouraged to participate and interact with their children throughout the session using the same materials. The clinician or researcher facilitating the training then has the opportunity to build on concepts or MRI-related stimuli previously introduced by parents in their home, and the bridged experience is less likely to be overly novel or intimidating for the child. Given that each child underwent mock training together with their co-twin, each participant also has to opportunity to observe their sibling who functions as a natural peer model. Systematic review findings suggest that sibling-involvement in ASD interventions perform generally similar to peer-mediated strategies, with positive outcomes in increased skill acquisition or reductions in unwanted behaviours (<u>Shivers & Plavnick, 2014</u>). Consistent with our protocol design, the effectiveness of sibling peer-modelling is further complemented by peer and parental prompting, the inclusion of direction instruction, and positive reinforcement strategies (<u>Watkins et al., 2014</u>). Where necessary, parents themselves may also act as a peer model by participating in the mock training procedure as their child observes the process.

### Conclusions and Future directions

In summary, core components implemented in the present MRI training protocol are as follows:

1. Pre-visit interview with parents to identify individualised reinforcers and preferred activities
2. Graduated exposure to stimuli encountered in scanner environment
3. Delivery of pre-visit scanner familiarisation materials by a familiar person
4. Peer-modelling with sibling-involvement
5. Minimize identified triggers or aversive events in the mock scanning environment
6. High level of child engagement with parental involvement and effective communication
7. Consistent and predictable scheduling, in particular with the use of visual schedules

Essential aspects of the MRI training procedure were the implementation of a child-focused approach, and effective familiarisation strategies specific to this population. While the sample size and absence of subjective outcome measures limit generalizability, an important consideration was to avoid unnecessary participant burden and fatigue. While including multiple measures of anxiety or adjustment levels pre- and post-training would ideally allow direct assessment of the efficacy of such strategies, the tradeoff would likely come at the cost of excessive demands on at-risk participants already engaged in a challenging procedure. This constrains selective and careful inclusion of measures and MRI sequences to those that are necessary and most relevant to avoid over-burdening each child. Depending on the research aims, repeating image acquisition sequences of interest may be of value. Given such limitations to inclusion of measures, decisions should therefore be carefully considered and evaluated with respect to the overarching aim and outcome of each research study.

Interestingly, we found some preliminary evidence that out-of-scanner head motion could predict that observed in-scanner during the MRI scan. This secondary finding is consistent with recent efforts to mitigate the effects of head motion with real-time feedback during mock training, where outcomes in head movement reduction were demonstrated to be generalizable to an actual MRI setting (<u>Cox, Virues-Ortega, Julio, & Martin, 2017</u>; <u>Greene et al., 2018</u>). The procedure involves training participants to reduce head movement by providing immediate visual feedback once head motion exceeds a certain threshold. Together with individualised familiarisation and reinforcement strategies, these findings are promising and extend current literature on improving image acquisition in challenging clinical populations such as ASD. The present study reports data quality metrics based on an MRI training protocol implemented for a local sample that was highly satisfactory compared to a large multi-site ASD cohort, and may be a valuable reference for future investigations on essential components for successful neuroimaging or similar task-related preparation.

## Acknowledgements

This research was conducted within the Developmental Imaging research group, Murdoch Children’s Research Institute and the Children’s MRI Centre, Royal Children’ s Hospital, Melbourne, Victoria. It was supported by the Murdoch Children’s Research Institute, the Royal Children’s Hospital, Department of Paediatrics The University of Melbourne and the Victorian Government’ s Operational Infrastructure Support Program. The project was generously supported by RCH1000, a unique arm of The Royal Children’s Hospital Foundation devoted to raising funds for research at The Royal Children’s Hospital. Research was conducted with support from Twins Research Australia. The authors declare no potential conflicts of interest with respect to the research, authorship, and/or publication of this article.

## Footnotes

1 https://www.rch.org.au/kidsinfo/fact_sheets/MRI_scans/

2 https://www.rch.org.au/okee/

3 http://fcon_1000.projects.nitrc.org/indi/abide/abide_II.html

