## Supplementary materials for "Individualised MRI training for paediatric neuroimaging in autism spectrum disorders: A child-focused approach"

### Supplementary Material

**Sample audio clips of MRI acoustic recordings** : <https://soundcloud.com/user-74029050>

**Visual schedule:** Photographs used to visually explain each stage of the appointment

1.
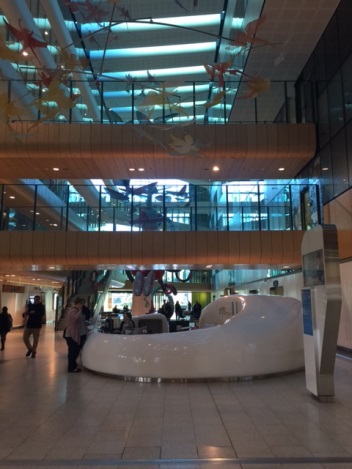
**Welcome to The Royal Children’s Hospital! (Main lobby to meet families)**
2.
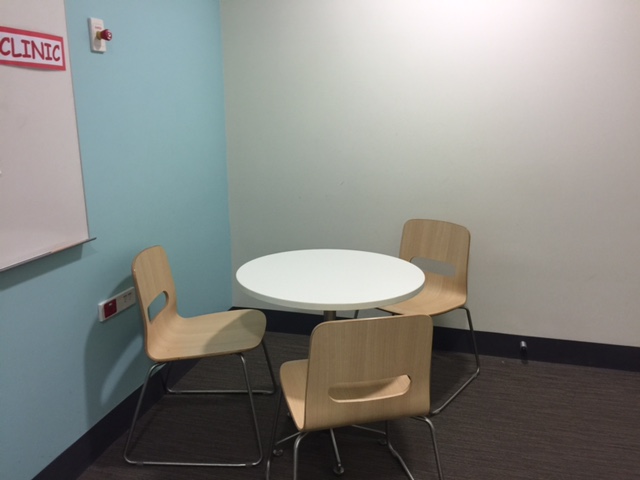
**Interview room for visit orientation**
3.
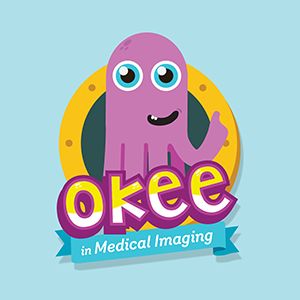
**Games from Okee mobile application, MRI orientation video**
4.
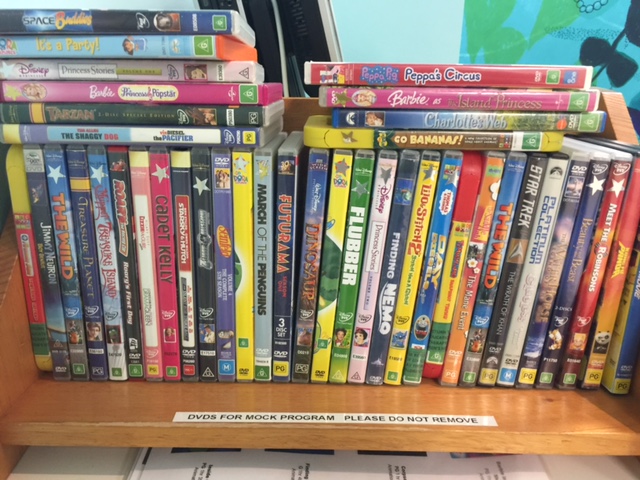
**DVD library selection for viewing in mock and actual MRI (or child’s own DVD)**
5.
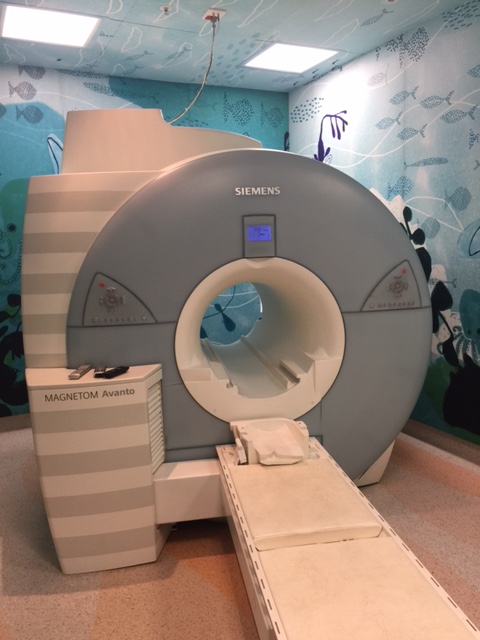
**Mock MRI scanner**
6.
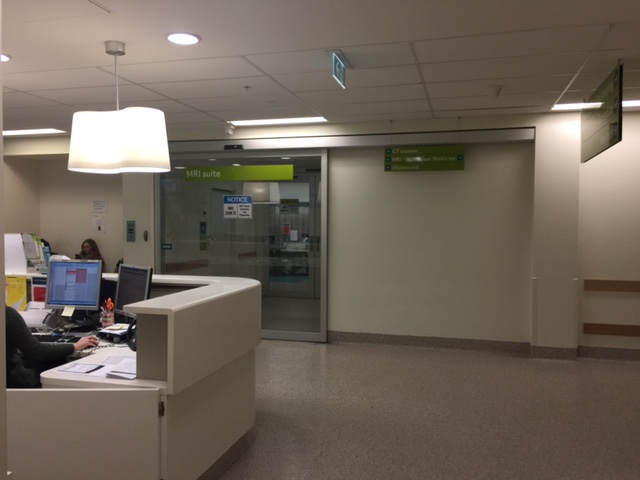
**RCH medical imaging reception**
